# AIDO.Tissue: Spatial Cell-Guided Pretraining for Scalable Spatial Transcriptomics Foundation Model

**DOI:** 10.1101/2025.07.04.663102

**Authors:** Jing Gong, Yixuan Wang, Nichloas Ho, Xingyi Cheng, Le Song, Eric Xing

**Affiliations:** GenBio AI; Mohamed bin Zayed University of Artificial Intelligence; Carnegie Mellon University

## Abstract

Single-cell spatial transcriptomics enables high-resolution insights into tissue organization and cell-cell interactions, yet poses significant computational and modeling challenges due to its scale and complexity. Here we introduce AIDO.Tissue, a spatially-informed pretraining framework. The design employs multiple cells as input and an asymmetric encoder-decoder architecture, making it effectively encodes cross-cell dependencies while scaling to large data. Systematic evaluation shows that our method scales with neighboring size and achieves state-of-the-art performance across diverse downstream tasks, including spatial cell type classification, cell niche type prediction and cell density estimation. These results highlight the importance of spatial context in building general-purpose foundation models for tissue-level understanding.

## 1. Introduction

Spatial transcriptomics technologies have enabled simultaneous measurement of gene expression and spatial coordinates across hundreds of thousands of cells, revealing critical spatial organization principles in diverse tissues (Marx, 2021; Moses & Pachter, 2022). These datasets capture essential cellular interactions, including cell-cell communication and spatial gradients that define tissue microenvironments (Fischer et al., 2023; Varrone et al., 2024). As spatial omics data continues to grow in scale, it presents an opportunity to learn spatially aware foundational representations of cellular variation.

Recent single-cell foundation models have demonstrated remarkable capabilities in learning generalizable representations through transformer-based architectures trained on tens of millions of cells (Theodoris et al., 2023; Cui et al., 2024; Hao et al., 2024; Kalfon et al., 2025). However, these models are pretrained on dissociated cell data without spatial context, limiting their ability to handle spatial transcriptomics tasks that depend on understanding cellular neighborhoods and tissue organization. Foundation models that have been exposed to both single-cell and spatial transcriptomics data have recently emerged, but existing spatial-aware models like CellPLM (Wen et al., 2023) and Nicheformer (Schaar et al., 2024) lack gene-level cross-cell attention. These methods fail to capture that cellular behavior is intrinsically linked to spatial context, given that cells respond to immediate neighbors and organize into tissue architectures that determine organ function (Lewis et al., 2021).

We introduce AIDO.Tissue, a novel spatial cell-guided pre-training framework tailored for foundation models in spatial transcriptomics. Our approach is built on two key innovations: (1) Explicit incorporation of spatial neighbor information—by taking multiple neighboring cells as input, the model learns both intra-cellular and inter-cellular dependencies, capturing richer spatial context; and (2) An asymmetrical encoder-decoder architecture—the encoder processes only the expressed genes across multiple cells, while the decoder focuses exclusively on reconstructing the gene expression of the center cell. This design significantly reduces computational overhead while maximizing the model’s ability to capture cross-cell, gene-level patterns critical for spatial representation learning.

Through systematic evaluation across two model scales (3M and 60M parameters) and multiple neighborhood sizes (8 to 64 cells), we demonstrate that spatial cell information is more important than scaling model size only. AIDO.Tissue also achieve a better performance than other competing method across diverse spatial related downstream tasks, including cell type classification, niche type prediction and cell density estimation. Our results suggest that incorporating spatial awareness during pretraining is crucial for building foundation models that can truly understand tissue biology, paving the way for more effective analysis of spatial transcriptomics data at scale.

## 2. Method

AIDO.Tissue incorporates spatial cell information to benefit pre-training large-scale single-cell RNA-seq data (illustrated in Figure 1). Instead of single individual cell as input, neighboring cells are also encoded simultaneously as joint input, which concept is analogous to MSA (Multiple Seuqence Alignment) style protein pretraining. The introduction of an asymmetrical encoder-decoder makes it computationally efficient to manipulate cross-cell gene dependency. We describe each component as below:

**Figure 1.**
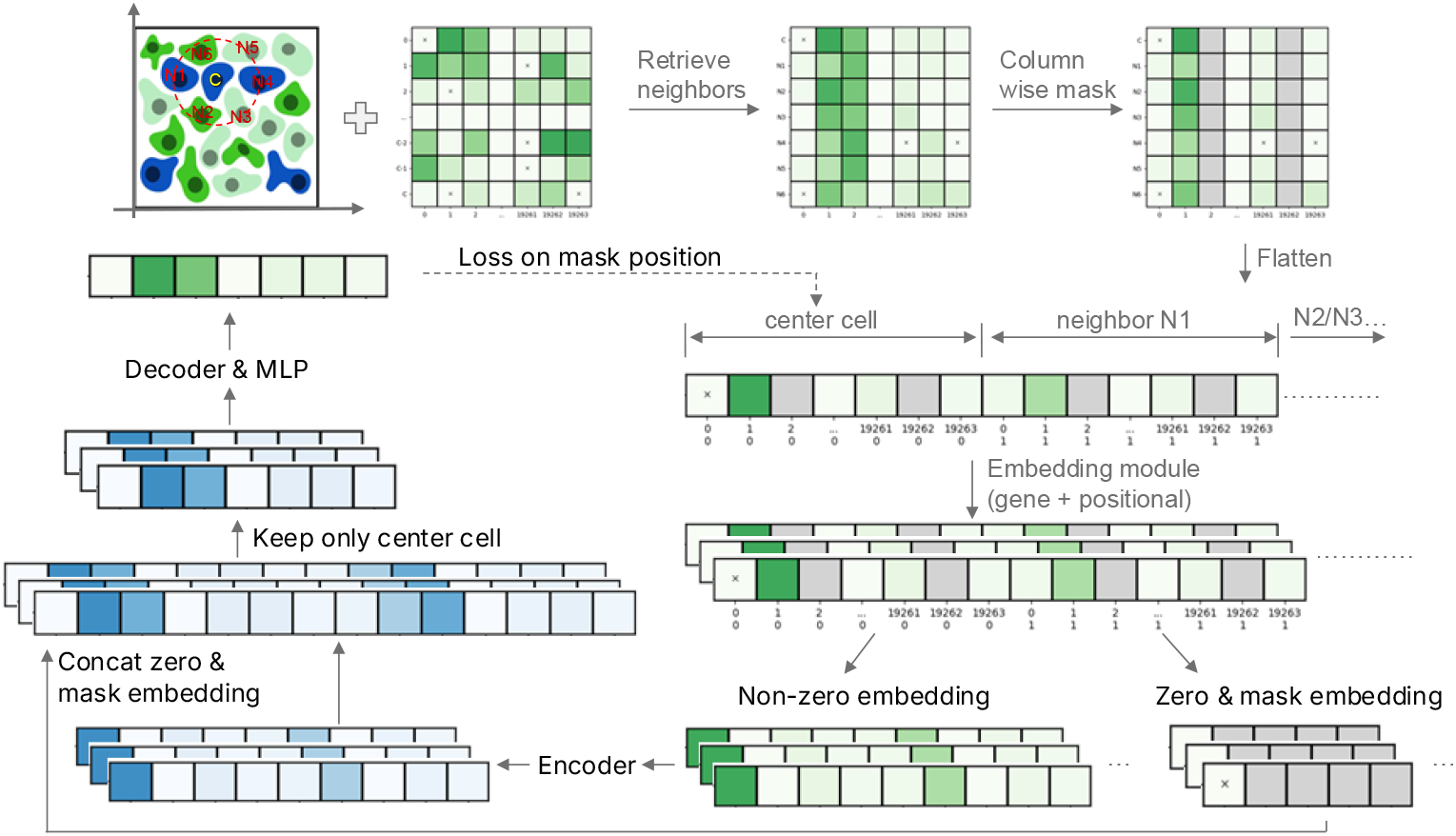
Overview of AIDO.Tissue spatial cell-guided pretraining architecture. The input is paired single-cell spatial and expression profiles. Each cell along with its retrieved *k* nearest neighbors are concatenated as a multi-cell input. The encoder processes expressed gene embeddings across all cells, while the decoder selectively reconstructs only the center cell. Please refer to the main text for a detailed explanation.

### Input

The input data consists of a paired single cell spatial profile matrix (*G* ∈ ℝ^*c×*2^) and expression matrix (*E* ∈ ℝ^*c×n*^), where *c* is number of cells and *n* is number of genes (in our study 19,264). The spatial matrix *G* denotes the x and y coordinate of each cell center in two-dimensional space. The expression matrix *E* contains normalized expression value of each gene across all cells, including expressed (non-zero count) and non-expressed (zero count) genes.

## Retrieving neighboring cells

For each cell, *k* ∈ (8, 16, 32, 64) nearest neighboring cells are retrieved based on cell-cell distance, which is calculated from spatial *G*. The expression vector of the center cell and neighboring cells is stacked into a larger matrix.

### Column masking

The overall pretraining objective is to recover masked expression values of the center cell. A column-wise masking strategy is introduced to avoid direct inference from the same gene of neighboring cells. The masking includes both non-zero and zero positions, but with a different ratio due to intrinsic abundance bias (see xTrimoGene (Gong et al., 2023) for more details).

### Flatten

To capture the dependency of genes between the center and neighboring cells, the masked matrix is converted to a longer flat one. A two-dimension vector is also employed to distinguish each gene and cell, where the first dimension is gene index and the second is cell index.

### Embedding

Each gene is encoded into a latent vector *d*, which is an element-wise sum of gene name embedding, expression value embedding and positional embedding. The gene name embedding is retrieved from an randomly initialized lookup embedding table. The expression value is projected to an embedding using an MLP-based module. Specifically, a rotary positional embedding is derived for each gene based on the two-dimensional gene and cell index.

### Encoder

For the full-length center-neighbor gene embedding, only expressed genes are kept and fed into the encoder. The design makes it efficient and computational affordable in Transformer-like architecture, especially when extending to a large number of neighboring cells. The attention mechanism of the encoder are calculated along all the input, thus capturing both inter-cell and intra-cell gene-gene dependency.

### Decoder

The output embedding of encoder contains latent information of all expressed genes. Before fed into decoder, the masked and non-expressed parts are also concatenated into a full-length one. To further reduce computational resources, only the center cell is fed into the decoder. Following the decoder, an MLP module is utilized to project the latent embedding into exact expression values.

### Loss calculation

The mean squared error (MSE) loss is employed to measure the error between the ground truth and the predicted expression value. The calculation is based on the masked positions of the center cell.

## 3. Experiments

### 3.1. Experimental Setup

#### Pretraining datasets

We collected large-scale spatial transcriptomics datasets across three main platforms for pertaining, including Vizgen (https://info.vizgen.com/ffpe-showcase), Nanostring (https://nanostring.com/resources) and 10xgenomics (https://www.10xgenomics.com/datasets). The final dataset contains about 76 slides and 22 million cells (See App.Table 1 for a detailed statistics.).

#### Pretraining configurations

The model was trained for a total of 150,000 iterations using a global batch size of 128. Optimization was performed using the Adam optimizer with *β*_1_ = 0.9 and *β*_2_ = 0.95, and a weight decay of 1 × 10^−2^ was applied to improve generalization. The learning rate was initialized at 2 ×10^−5^ and then warm-up to a 2 ×10^−4^ and then following a cosine decay schedule. To stabilize training, gradient clipping was employed with a maximum norm of 1.0. We pretrained 3M and 60M parameter (App.Table 2) transformer models with varying spatial neighborhood sizes *k* ∈ {8, 16, 32, 64}.

### 3.2. Scaling Law Analysis

To systematically evaluate the performance and capacity of our spatial pretraining framework, we first conducted a scaling analysis along the neighborhood size. Specifically, we varied the number of spatial neighboring cells incorporated into the input context to assess how much spatial information is necessary or beneficial. This neighbor size scaling provides insight into the locality of spatial gene expression patterns and the extent of spatial dependency learned by the model. Here we fine-tuned the model on the niche label prediction dataset as the benchmark.

As show in Figure 2, we observed a consistent improvement while increasing neighbor size from 8 to 64, suggesting that more neighboring cells provide a richer spatial context. The phenomenon is similar for the larger 60M parameter size model (see App. Figure 6). These scaling behavior demonstrates the effectiveness of our spatial cell-guided pretraining approach.

**Figure 2.**
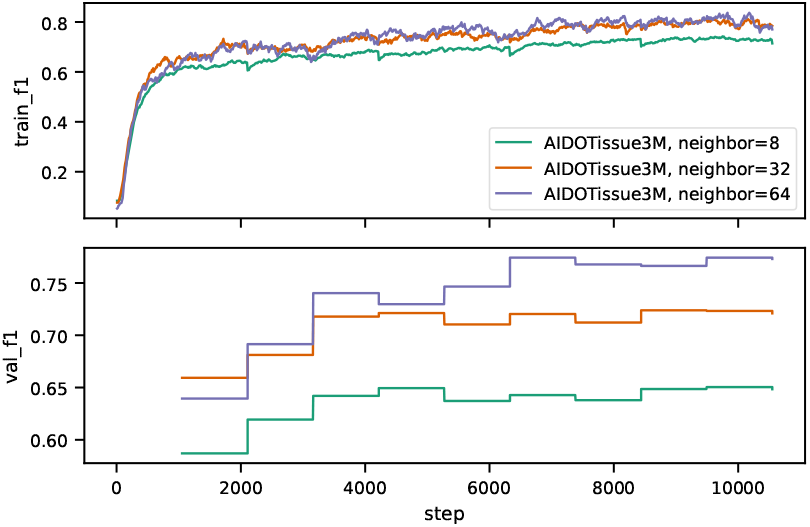
Fine-tuning metric curves for niche label prediction across different neighboring size configurations. Evaluated on 3M parameter size model.

However, we found only marginal improvement while further scaling the model size (32 neighbors 3M versus 60M, App. Figure 6, 7). The benefit is much smaller than that of increasing neighbor size (3M model from 32 neighbors to 64, App. Figure 6, 7). This indicates that, in spatial transcriptomic modeling, the bottleneck may not lie in model expressiveness but rather in the richness of the spatial information available.

### 3.3. Downstream Task Evaluation

To comprehensively assess the utility of our spatially pre-trained model, we benchmarked three downstream tasks that have been established in the NichFormer framework. We use the CosMx human liver dataset from CosMx data resource (He et al., 2021) for cell type and niche type benchmarking and Xenium human lung dataset from the 10x Genomics data resource for cell density evaluation. In the following sections, we detail the definition and results for each task, highlighting the model’s performance and behavior relative to existing baselines.

#### 3.3.1. Cell Type Prediction

This classification task involves assigning one of 22 well-annotated cell types to each cell based on both its gene expression and spatial context. Unlike traditional cell type annotation tasks that rely solely on transcriptomic profiles, this dataset also provides spatial information.

We observed that AIDO.Tissue outperforms Nicheformer in prediction (F1 score 0.77 versus 0.73, Figure 3 (A)), highlighting the advantage of incorporating richer spatial context. While Nicheformer leverages both single-cell and spatial transcriptomics data during pretraining, its architecture processes each sample centered on a single cell, limiting the spatial information to what can be implicitly learned from pairwise representations. In contrast, AIDO.Tissue explicitly integrates the gene expression and spatial embeddings of neighboring cells during both pretraining and inference. This design enables the model to capture local tissue structure and microenvironmental signals more effectively, leading to improved cell identity resolution.

**Figure 3.**
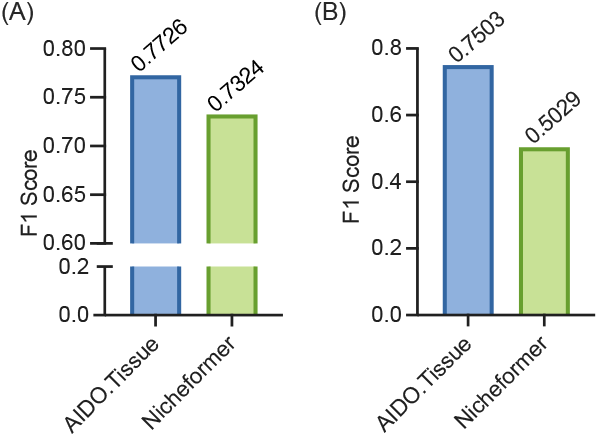
Cell type (A) and niche type (B) prediction performance comparison with F1 score metric.

#### 3.3.2. Niche Type Prediction

Following spatial cell type classification dataset, there derives a microenvironment-level prediction task: niche type prediction task. The task focuses on classifying each cell into one of 6 predefined spatial niches, which are aggregated from 22 original cell types based on shared spatial localization and functional roles. These niche types represent coherent microenvironmental structures, such as immune-rich regions, stromal domains, or tumor cores, and serve as a biologically meaningful abstraction that captures both cellular identity and spatial context.

We found that our approach achieves substantial improvements over baseline Nicheformer, with F1-scores from 0.50 to 0.75 (Figure 3 (B)). We also visualize one region with ground truth and predicted niche types. It shows our prediction has a clear delineation of tissue regions and smooth transitions between niche types, which is well aligned with the truth labels (Figure 4, App. Figure 8). These results highlight the model’s ability not only to classify cells accurately but also to infer higher-order spatial organization, demonstrating its potential utility in both diagnostic and discovery-oriented spatial omics applications.

**Figure 4.**
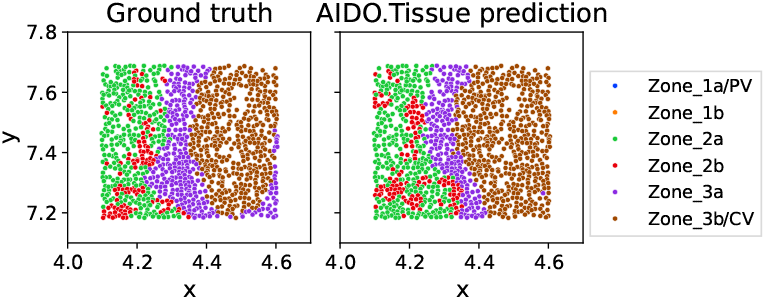
Visualization of the ground truth niche type and predicted niche type distribution of a region from test set.

#### 3.3.3. Cell Density Prediction

This regression-based task aims to estimate the local cellular composition around a given center cell by predicting the proportion of each cell type within a defined spatial radius. Such local density distributions often reflect tissue organization and microenvironmental context and are known to differ substantially between healthy and tumor tissues. We use MAE (Mean Absolute Error) as the evaluated metric.

As shown in Figure 5, the AIDO.Tissue model achieves a lower MAE value (4.583) than Nicheformer (7.084), indicating its superior capacity to reconstruct fine-scale cellular distributions (see App. Figure 9 for more metrics). This improvement suggests that our model can integrate broader neighborhood context effectively.

**Figure 5.**
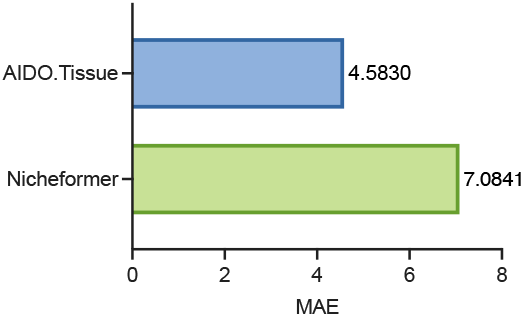
Cell density prediction performance measured by MAE.

## 4. Conclusion

We introduce AIDO.Tissue, a novel and efficient framework to pretrain transcriptomic data in a spatial cell-guided manner. Through systematic scaling analysis, we demonstrate that spatial neighborhood size often has greater impact on downstream performance than raw model capacity. By integrating spatial neighboring cell information, we observed an consistent improvement across diverse downstream tasks, which illustrates that spatial context is a fundamental organizing principle that should be incorporated during the pretraining phase. The AIDO.Tissue framework provides a scalable foundation for analyzing increasingly complex spatial transcriptomics datasets, paving the way for deeper understanding of tissue organization and cellular interactions at unprecedented scale. Code and pretrained model weights are publicly available at https://github.com/genbio-ai/ModelGenerator/tree/main.

## A. Appendix materials

**Table 1.** Statistics of pretraining single-cell spatial data.

| Platform | Number of slides | Number of cells (million) |
| --- | --- | --- |
| Vizgen | 19 | 9.3 |
| Nanostring | 12 | 1.7 |
| 10xgenomics | 45 | 10.7 |
| Total | 76 | 21.7 |

**Table 2.** Hyper-parameters of the pre-trained models.

| Model name | Parameter (M) | Encoder depth | Encoder heads | Encoder dim | Decoder depth | Decoder heads | Decoder dim | Neighbor number |
| --- | --- | --- | --- | --- | --- | --- | --- | --- |
| AIDO.Tissue-3M | 3 | 4 | 2 | 128 | 2 | 2 | 128 | 8, 32, 64 |
| AIDO.Tissue-60M | 60 | 12 | 8 | 512 | 4 | 8 | 512 | 16, 32 |

**Figure 6.**
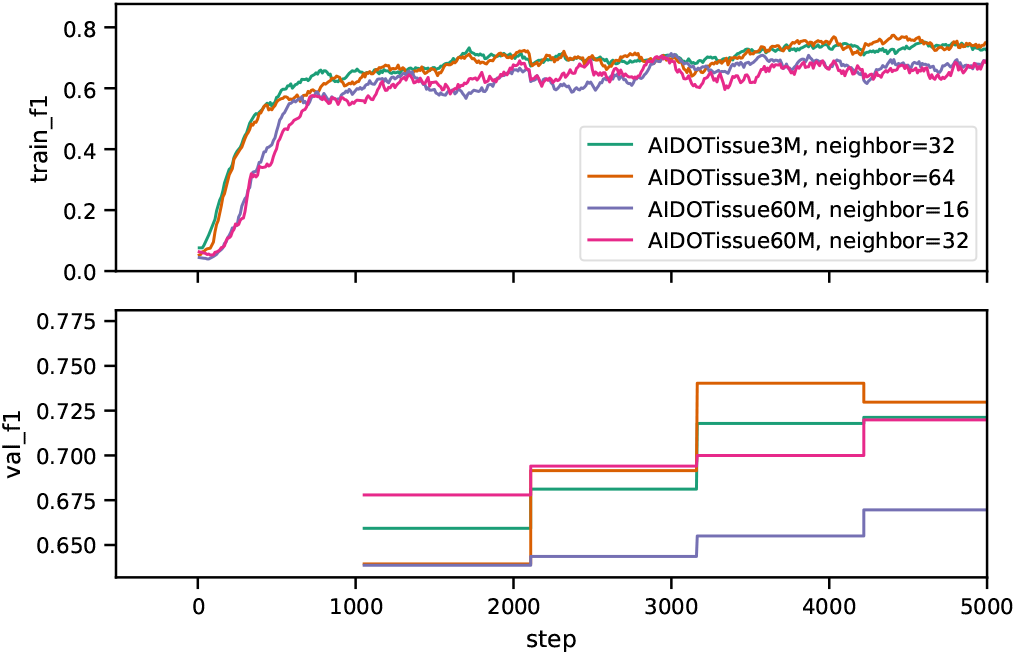
Fine-tuning metric curves for niche label prediction across different model size configurations.

**Figure 7.**
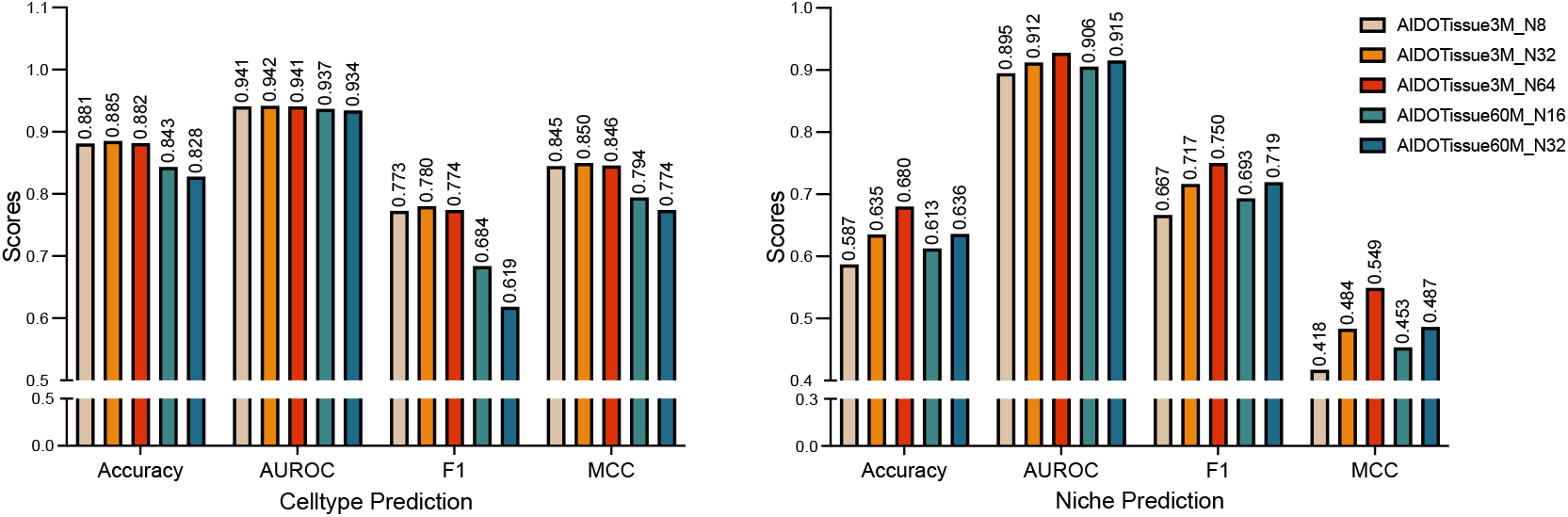
Performance comparison on cell type (left panel) and niche type prediction (right panel) across different model configurations.

**Figure 8.**
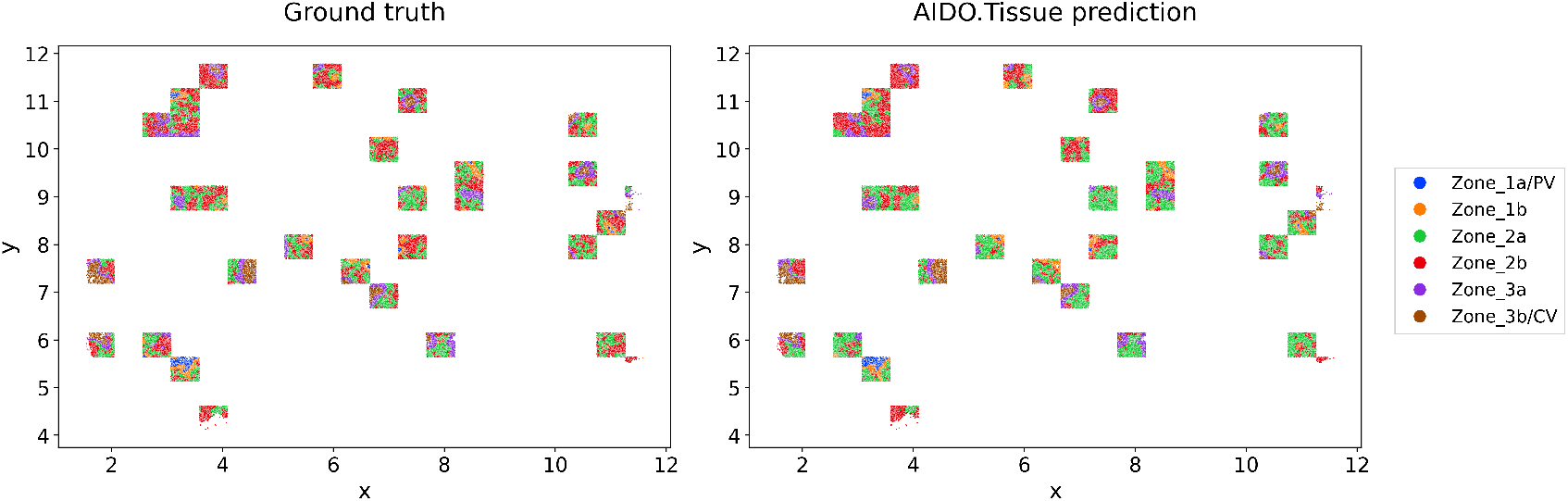
Full visualization of the ground truth niche type and predicted niche type distribution. All the test set regions are plotted.

**Figure 9.**
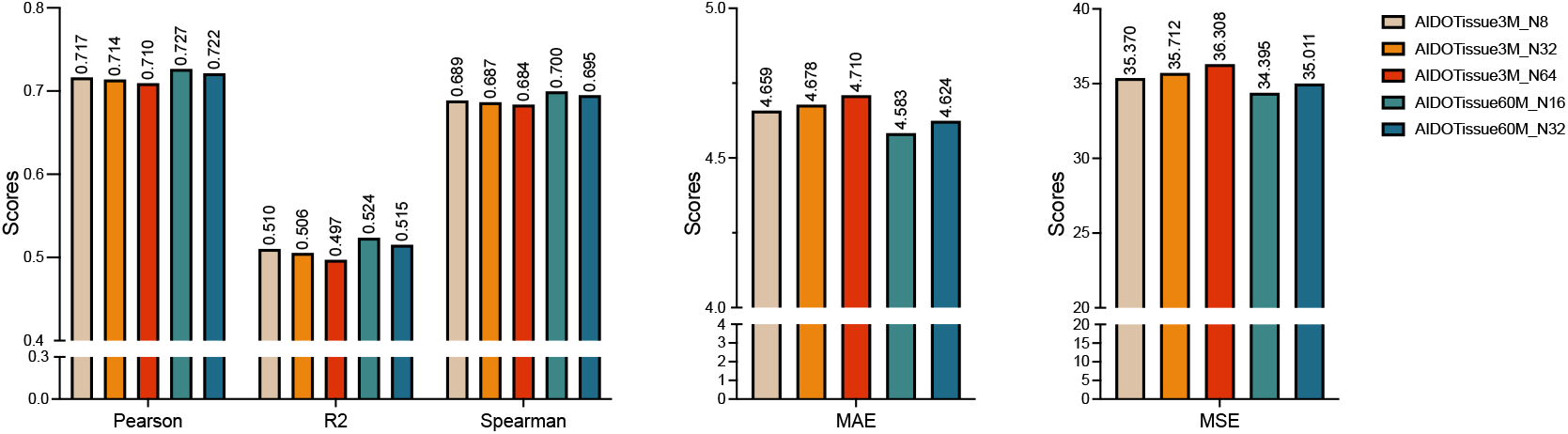
Performance comparison on cell density prediction across different model configurations.

